# Meiotic DSB repair DNA synthesis tracts in *Arabidopsis thaliana*

**DOI:** 10.1101/2024.02.25.582015

**Authors:** Miguel Hernández Sánchez-Rebato, Veit Schubert, Charles I. White

## Abstract

We report here the successful labelling of meiotic prophase I DNA synthesis in the flowering plant, *Arabidopsis thaliana*. Incorporation of the thymidine analogue, EdU, enables visualisation of the footprints of recombinational repair of programmed meiotic DNA double-strand breaks (DSB), with ∼400 discrete, SPO11-dependent, EdU-labelled chromosomal foci clearly visible at pachytene and later stages of meiosis. This number equates well with previous estimations of 200-300 DNA double-strand breaks per meiosis in Arabidopsis, confirming the power of this approach to detect the repair of most or all SPO11-dependent meiotic DSB repair recombination. The chromosomal distribution of these DNA-synthesis foci accords with that of early recombination markers and MLH1, which marks Class I crossover sites, colocalises with the EdU foci.

It is currently estimated that ∼10 cross-overs (CO) and an equivalent number of non-cross-overs (NCO) occur in each Arabidopsis male meiosis. Thus, at least 90% of meiotic recombination events, and very probably more, have not previously been accessible for analysis. Visual examination of the patterns of the foci on the synapsed pachytene chromosomes corresponds well with expectations from the different mechanisms of meiotic recombination and notably, no evidence for long Break-Induced Replication DNA synthesis tracts was found. Labelling of meiotic prophase I, SPO11-dependent DNA synthesis holds great promise for further understanding of the molecular mechanisms of meiotic recombination, at the heart of reproduction and evolution of eukaryotes.

**Author Summary:** Sexual reproduction involves the fusion of two cells, one from each parent. To maintain a stable chromosome complement across generations, these specialized reproductive cells must be produced through a specialized cell division called meiosis. Meiosis halves the chromosome complement of gametes and recombines the parental genetic contributions in each gamete, generating the genetic variation that drives evolution. The complex mechanisms of meiotic recombination have been intensely studied for many years and we now know that it involves the repair of programmed chromosomal breaks through recombination with intact template DNA sequences on another chromatid. At the molecular level, this is known to involve new DNA synthesis at the sites of repair/recombination and we report here the successful identification and characterisation of this DNA neo-synthesis during meiosis in the flowering plant, Arabidopsis. Both the characteristics and numbers of these DNA synthesis tracts accord with expectations from theory and earlier studies. Potentially applicable to studies in many organisms, this approach provides indelible footprints in the chromosomes and has the great advantage of freeing researchers from dependence on indirect methods involving detection of proteins involved in these dynamic processes.

## Introduction

Meiosis is the specialised cell division that halves chromosome numbers during the production of gametes, essential for the maintenance of stable ploidy across generations in sexually-reproducing organisms. In contrast to mitosis, in which duplication of the genome is followed by a single cellular division, meiosis involves one round of DNA replication followed by two rounds of division. Separating sister chromatids to opposite poles, the second meiotic division is, in broad terms, analogous to mitosis. Balanced chromosomal segregation during the first division however requires that homologous chromosomes recognise each other, pair and orient themselves on the spindle so that the two members of each pair move to opposite poles during anaphase. This is ensured by the programmed induction of homologous recombination through the induction and repair of DNA double-strand breaks (DSB) during the first meiotic prophase.

This programmed, highly regulated process of DNA damage and repair is essential for proper meiotic chromosome alignment and segregation and thus fertility in eukaryotes, with some known exceptions, notably *Drosophila* and *C. elegans* (see review by [1]). Following the programmed induction of DSB by the SPO11 complex, the cleaved DNA ends are resected to produce 3’-ssDNA overhangs, onto which the recombinases DMC1 and RAD51 are loaded to form presynaptic nucleoprotein filaments. These presynaptic nucleofilaments are the active molecular species in the search for, and invasion of an intact homologous DNA template, the first steps of DNA repair via homologous recombination. DNA synthesis then extends the invading 3’ end across the template and the joint-molecule intermediate structures are further processed to yield the two main products of HR-mediated DSB repair: gene conversions associated with reciprocal exchange of their flanking sequences (crossovers, CO), or not (non-crossovers, NCO).

DNA synthesis is thus inherent to DSB repair via homologous recombination. In the 1960s, studies based on incorporation of H3-thymidine and C14-thymidine into newly synthesised DNA gave evidence for meiotic prophase I DNA synthesis in the plants *Lilium longiflorum* and *Trillium erectum*, the amphibian *Triturum viridescens*, mice and humans [2–6]. These early studies have since been extended to show association of DNA synthesis with pachytene recombination nodules in *Drosophila* female meiosis and with the synaptonemal complex in mouse spermatocytes [7,8].

Labelling with 5-bromo-2’ -deoxyuridine (BrdU) in budding yeast has been used to confirm the presence of SPO11-dependent meiotic prophase DNA synthesis [9]. The authors characterized both single-molecule BrdU tracts at a recombination hotspot and the timing of their appearance in relation to other meiotic features. Observation of SPO11-dependent, BrdU labelling pattern has also been described in *Tetrahymena* [10] but, to our knowledge, no further data has been published in other organisms. In Arabidopsis, incorporation of BrdU or 5-ethynyl-2’-deoxyuridine (EdU) during pre-meiotic S-phase has been used to establish the meiotic timeline of meiosis in Arabidopsis pollen mother cells (PMC) [11–13]. There are thus far no reports of labelling non-S-Phase DNA synthesis during Arabidopsis meiosis.

The induction of synchronous meiosis in populations of cells has been key to a number of major advances in understanding of the molecular processes that make up the meiotic division [14–18]. However, such direct approaches have not been available in other organisms and this has resulted in significant gaps in understanding. A clear example of this is seen in the flowering plant, *A. thaliana*. Estimations of numbers of SPO11-induced DSB in Arabidopsis pollen mother cells (PMC, male meiosis) are ∼200-300 per meiosis. Of these, ∼10 yield inter-homologue CO and another 10 have been shown to give inter-homologue NCO [19,20]. Thus, the repair outcome of at least 90% of meiotic DSB in Arabidopsis is not known. The situation is strikingly different in budding yeast with 66 NCO plus 90.5 CO estimated per meiosis, the sum of which is very close to the estimated 140-170 meiotic DSB. Similarly in mouse, 273 NCO plus 27 CO sum close to the estimated 200-400 meiotic DSB [21–24].

Detection of recombination-associated DNA synthesis tracts offers an attractive path to understand the mechanisms underlying these “invisible” meiotic recombination events in Arabidopsis and potentially in other organisms. We present here the successful labelling of SPO11-dependent, meiotic prophase I DNA synthesis by EdU incorporation in Arabidopsis PMC.

## Results

### EdU labelling of DSB repair-associated DNA synthesis

In Arabidopsis, while each inflorescence contains flower buds bearing PMC at different pre-meiotic and meiotic stages, the PMC of the anthers of a given flower bud are nearly synchronous [25,26]. Along with the sequence of cytologically distinguishable meiotic stages in the buds of a given inflorescence, in practice it is relatively straightforward to classify PMC subpopulations within the meiotic timeline.

Timelines of Arabidopsis male meiosis concord that pre-meiotic G2 phase lasts 7-10 hours and that pollen mother cells advance from the beginning of leptotene to the end of pachytene in 19 to 21 hours [11–13,27](summarized in Figure 1). Subsequent stages of meiosis are considerably faster, with post-pachytene meiosis estimated to take from 2.7 to 9 hours. Given that meiotic DSB formation begins at the onset of prophase I (early leptotene) and continues for a few hours [28], any meiotic prophase I DSB-repair DNA synthesis should occur at least 7-10 hours after the end of pre-meiotic S-phase. We thus decided to incubate the inflorescences with the nucleoside analogue for a 24-hour pulse to entirely cover the prophase I DSB repair-associated DNA synthesis window. Given the relative synchrony of progression through meiosis of the PMC from a given flower bud, the easily identifiable appearance of S-phase labelling and the possibility to easily situate subpopulations of meiotic cells in the temporal meiotic sequence, it seemed reasonable to expect that we would be able to exclude S-phase labelled meioses, and thus focus our analyses on those labelled during Prophase I.

**Figure 1.**
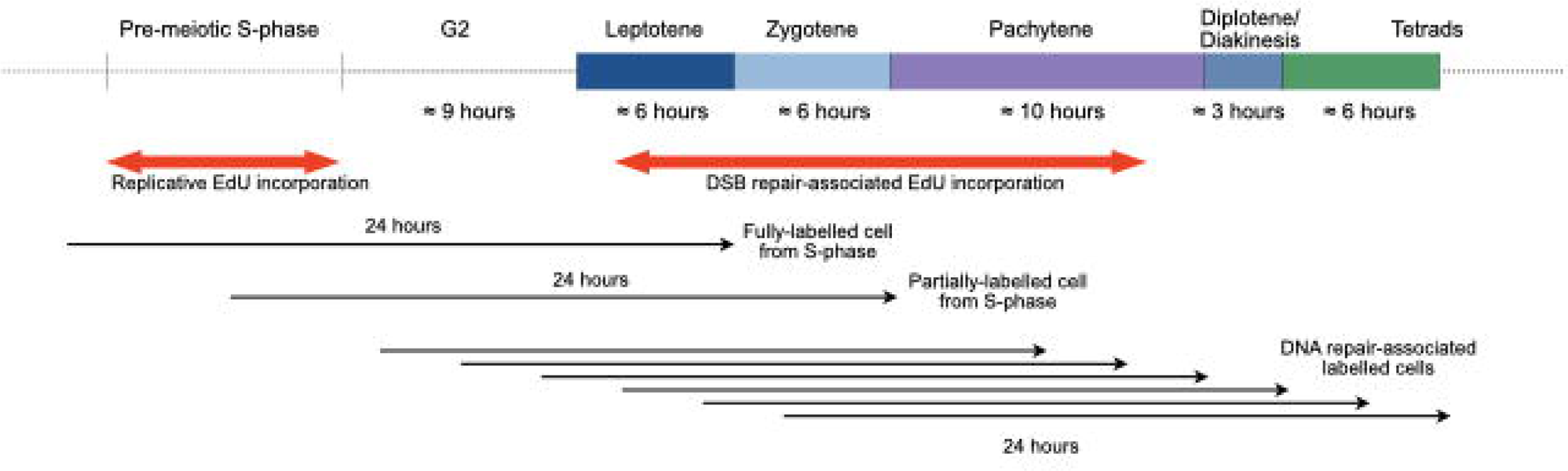
EdU labelling strategy and timeline of Arabidopsis male meiosis. Pollen mother cell nuclei were fixed and isolated at the end of a 24-hour pulse of EdU incorporporation. Cells which traversed all or part of the pre-meiotic S-phase during the pulse will show replicative EdU incorporation and be at G2/leptotene/zygotene stages at the end of the pulse, with some overlap into early pachytene. Fully pachytene nuclei (mid-late pachytene) will not show replicative labeling, but will have traversed the leptotene and zygotene stages in the presence of EdU. Any EdU incorporation in these (or later) nuclei will be due to non-S-Phase DNA synthesis.

Notwithstanding the current absence of direct evidence of the timing of occurrence of meiotic recombination intermediates in Arabidopsis, DAPI staining and immunolocalisation of meiotic proteins implicated in early invasion intermediates (RAD51/DMC1), joint molecules (MSH4, early HEI10) and resolution (MLH1, late HEI10) give a clear picture of the progression of key events through meiotic prophase I. In pachytene, when synapsis is complete, mean numbers of MLH1 foci correspond to the number of Class I crossovers (∼90% of total CO), measured by counting of chiasmata at metaphase I. In analogy to organisms for which recombination intermediates can be tracked directly, it is reasonable to assume that most recombination events are either resolved or close to resolution at pachytene in Arabidopsis.

As summarised in Figure 1, inflorescences were thus incubated with EdU for 24 hours to ensure coverage of the window of DSB repair-associated DNA synthesis through prophase I up to the end of pachytene. Pachytene cells observed at the end of the 24-hour pulse would have traversed prophase I up to that point in the presence of the analogue. As mentioned above, nuclei with pre-meiotic S-phase labelling were easily identified and excluded from the analysis.

### Non-replicative DNA synthesis tracts are present from pachytene until the end of meiosis

Early prophase I meiocytes, including cells in leptotene and zygotene, showed complete or extensive EdU labelling of their chromosomes (Figure 2a). This replicative labelling confirms, as expected, that they had passed through pre-meiotic S-Phase during the EdU pulse [11–13,29,30]. Furthermore, nuclei with partial replicative labelling of DAPI-dense, late replicating heterochromatic regions were observed, as expected for cells that were in mid/late S-phase at the beginning of the EdU pulse (second row of Figure 2a) [30–33].

**Figure 2.**
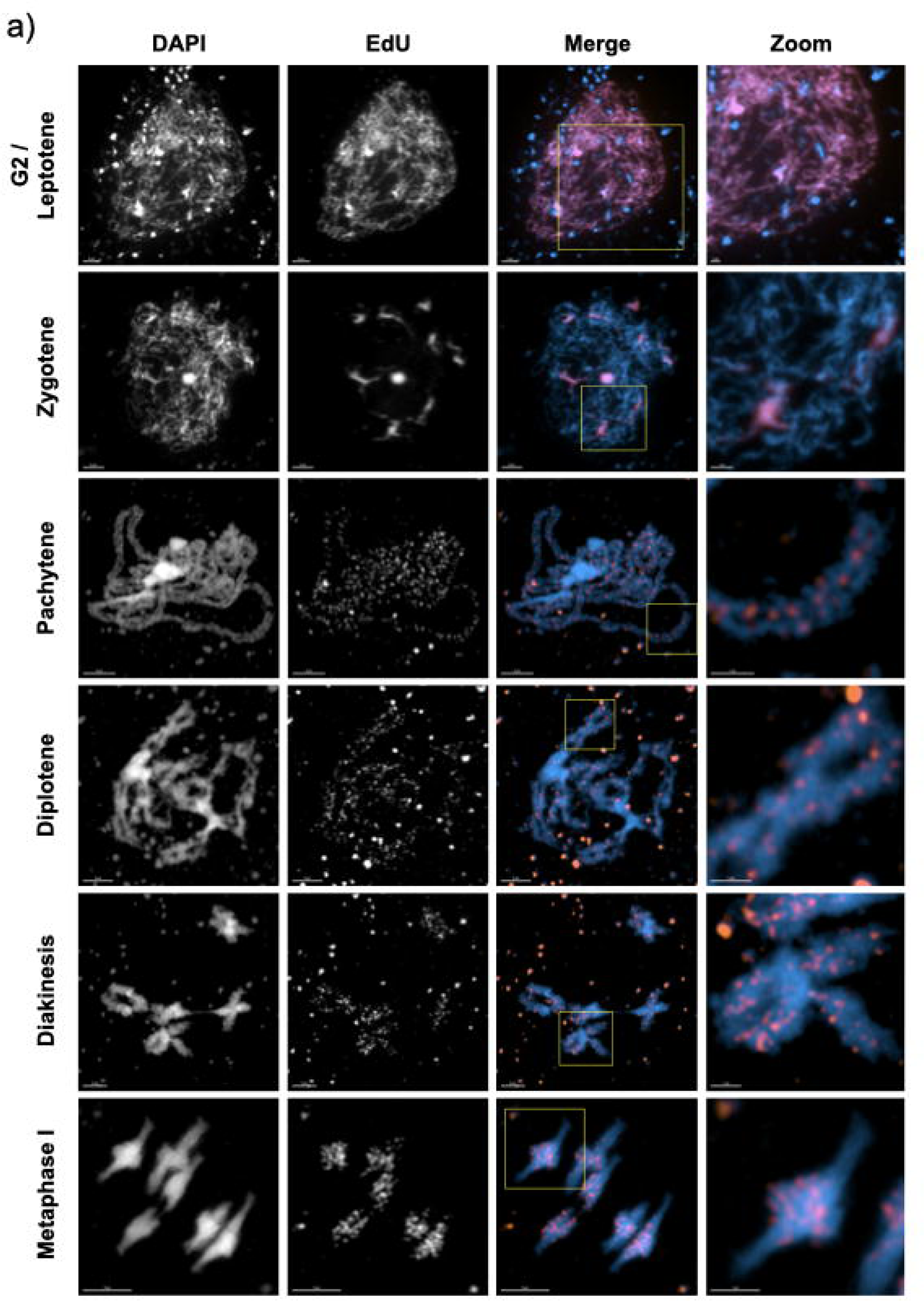
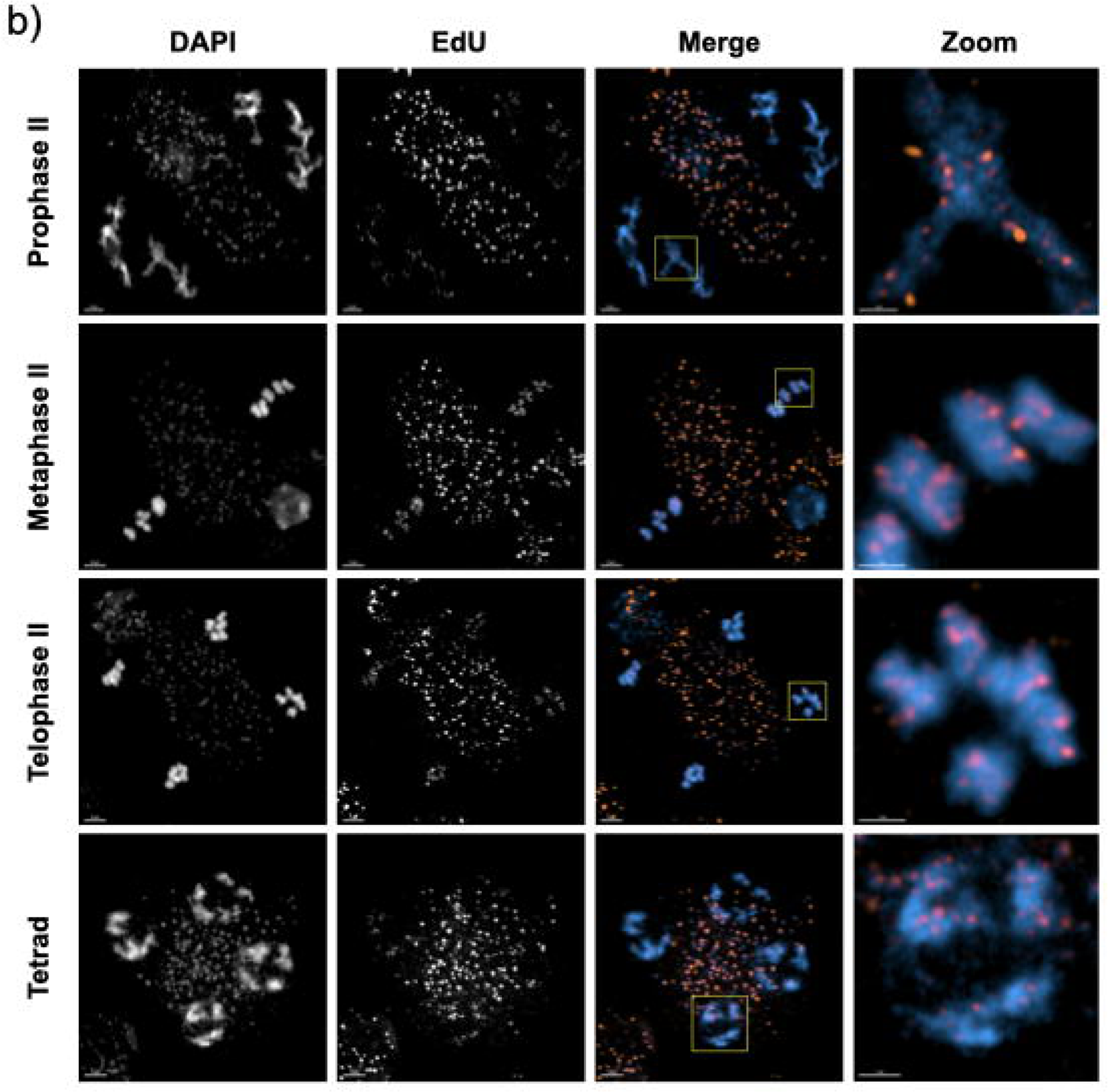
DSB repair-associated DNA synthesis tracts are detected from pachytene up to the end of meiosis. a) Meiosis I and b) meiosis II stages in *A. thaliana* wildtype. S-phase, replicative DNA labelling is seen at early G2/letotene and zygotene stages, while clear, discrete foci of EdU substituted DNA are visible in the chromosomes at later stages from pachytene onwards. Rectangles indicate the enlarged regions presented in the zooms. Non-chromosomal EdU foci correspond to labelling of mitochondrial genome replication. 3µm scale bars are shown at the bottom left of each image (1μm for the zooms). Images were taken with the confocal microscope + Airyscan module.

In striking contrast, pachytene nuclei showed faint, discrete foci of a considerably smaller size (Figure 2a Pachytene, see also Figure 4a and Figure S1). These foci are very faint through the microscope eyepiece, but are clearly visible when the images are processed. This difference in labelling intensity between S-phase and prophase EdU incorporation is not unexpected, considering that the latter would correspond to DNA synthesis tracts of tens to a few thousands of base pairs, while replicative DNA synthesis would range from megabase-long late-replicating regions to the whole genome (4C ≈ 520 Mb in these prophase I nuclei) [34]. We note also the presence of labelled mitochondria in these images, confirming their replication during pre-meiotic G2 and/or meiotic prophase I cells. These labelled mitochondria, clearly visible from pachytene on, are also present at earlier stages, but are not readily visible due to the very strong fluorescence of the S-phase replicative nuclear labelling (Figure 2).

The EdU foci observed in pachytene cells appear well-distributed across the chromosomes, but are less present in DAPI-dense, heterochromatic regions (Figure 2a Pachytene, Figure 4a). Furthermore, given the tighter-packed DNA in these regions, the observed lesser density of EdU foci is presumably an underestimation. This pattern of foci concords with genetic crossover mapping, studies of meiotic DSB (SPO11-oligo sequencing) and immunofluorescence studies of the distributions of early recombination proteins, all of which show meiotic recombination to follow a relatively even distribution across the Arabidopsis genome, with the exception of (heterochromatic) centromeric regions which have significantly lower densities of SPO11-oligos and crossovers [34–36].

As summarised in Figure 1, based on published timelines [11–13,27] and our own measurements, a meiocyte observed at the telophase II/tetrad stage would have been at leptotene/zygotene at the beginning of the 24-hour EdU pulse. We had thus good reason to expect that prophase I EdU foci would be observable in all meiotic stages up to the end of meiosis. This was indeed the case, with EdU foci visible from pachytene to the end of meiosis (telophase II/tetrad Figure 2). Meiotic chromosomes condense progressively as they traverse prophase I, evolving from a mass of hardly-distinguishable thin fibres at early leptotene to thick synapsed fibres in pachytene and to individualised dense bivalents with a more polygonal shape at metaphase I. This condensation produces significant shortening of the chromosomes and results in changes of the visual character of the EdU signals. While most EdU foci at pachytene appear as discrete spots, they merge into larger patches at diplotene and this continues through diakinesis, up to the greatest condensation at metaphase I (Figure 2, diplotene-metaphase I). Furthermore, heterochromatic, centromeric regions become more easily identifiable when bivalents individualize at diakinesis and the relatively low density of EdU signals in these regions becomes clearly visible. This is even more pronounced at metaphase I, with clear visual differentiation between the centromeric and pericentromeric regions of the two homologous chromosomes of each bivalent. Finally, the EdU signals are observed throughout the second meiotic division, changing their appearance with evolving chromosome condensation and shape in a way similar to that seen during the first meiotic division (Figure 2, prophase II-tetrad).

### Prophase I DNA synthesis is SPO11-dependent

Meiotic recombination is initiated by the induction of DNA DSB by the SPO11 complex at the beginning of meiotic prophase I. To verify that the discrete non-replicative DNA synthesis tracts observed at pachytene and subsequent stages originate from SPO11-dependent meiotic DSB repair, we carried out this analysis in parallel in *spo11-1* mutant plants [37] (Figure 3). Pachytene images were analysed with the IMARIS software package (Bitplane AG, Oxford Instruments, UK) *Spots* function. EdU foci were selected based on diameter and mean intensity in the EdU channel, and filtered to retain only those colocalising with chromosome fibres (mean intensity cutoff in the DAPI channel). A high-cut filter of mean intensity in the EdU channel was also applied to filter out the much-brighter organellar foci.

**Figure 3.**
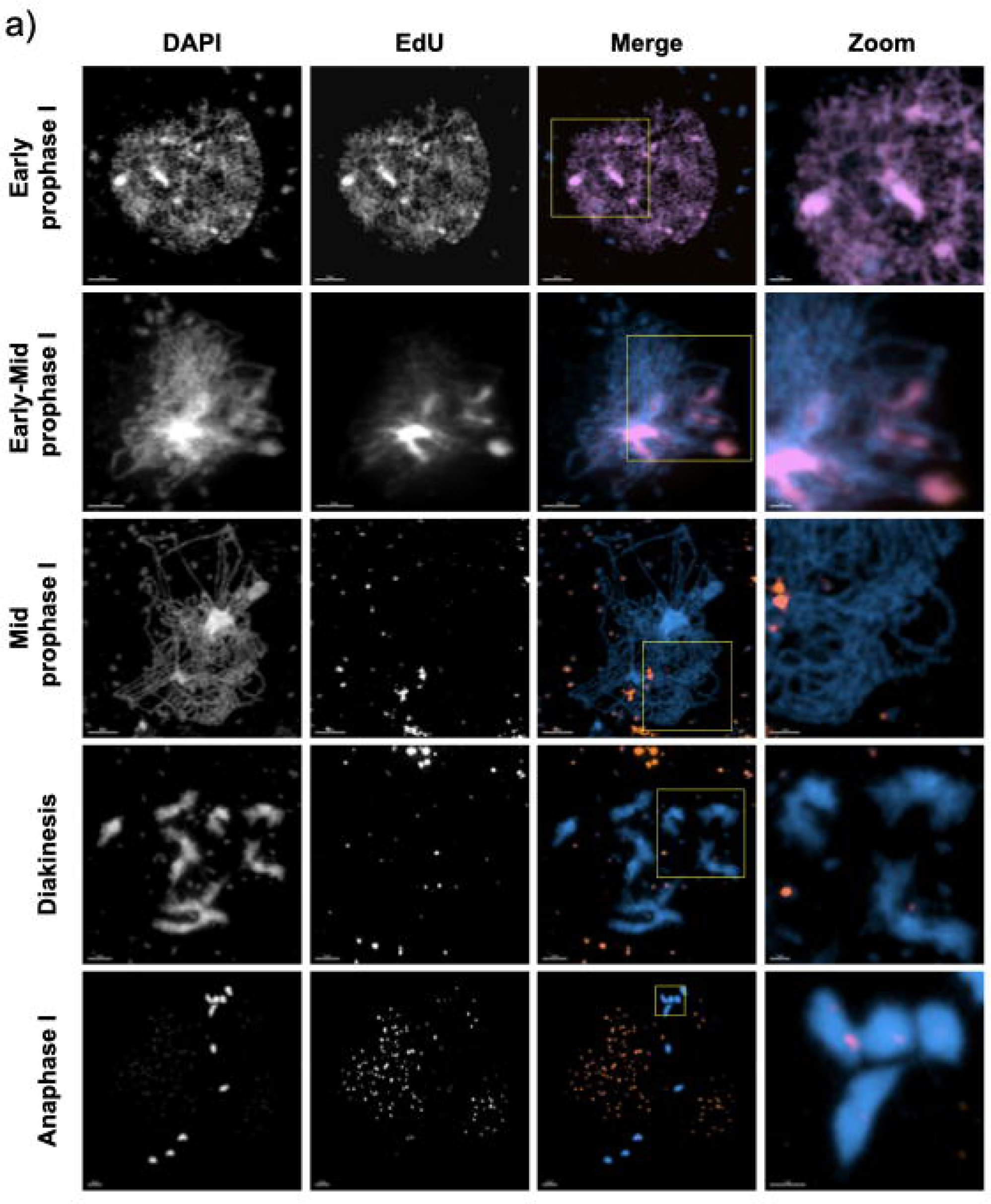
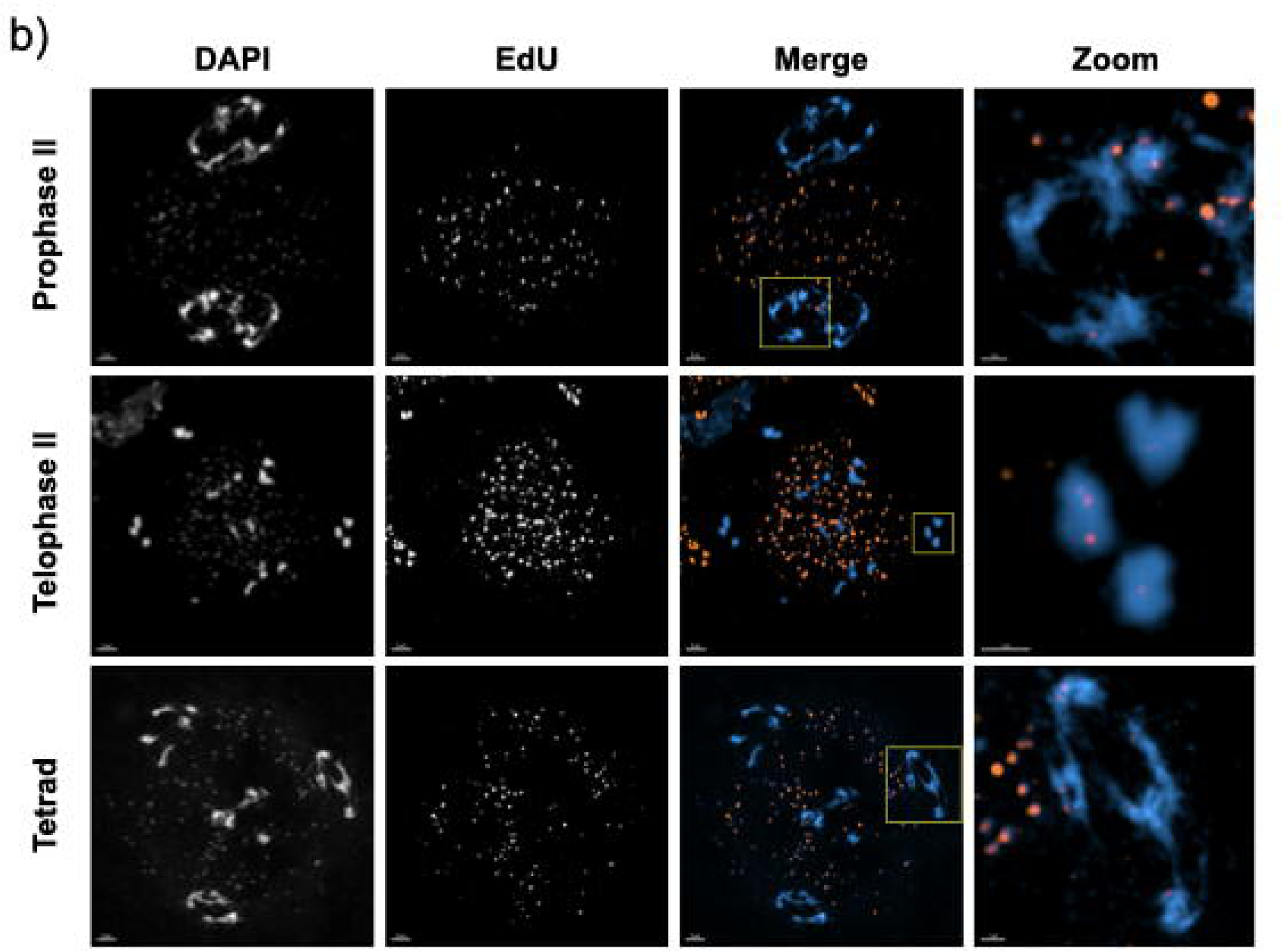
DSB repair-associated DNA synthesis is SPO11- dependent. a) Meiosis I and b) meiosis II stages in *A. thaliana* spo11-1 mutants. As for WT meiosis (Figure 2), replicative S-phase labelling is seen at early stages, corresponding to G2/leptotene and zygotene. As expected, a striking reduction in numbers of EdU foci (compared to WT) is observed in spo11-1 mutants at stages from mid-prophase I onwards (corresponding to pachytene and later stages). Rectangles indicate the enlarged regions presented in the zooms. Images taken with the confocal microscope with Airyscan module. 3µm scale bars are shown at the bottom left of each image (1μm for the zooms).

WT pachytenes show 409.5 ± 7.96 EdU foci per nucleus (mean ± SEM; n = 45 nuclei; Figure 2, Figure S1), with counts ranging from 323 to 510 (plus one outlier with 263). Ten of the 45 image stacks analysed were acquired via 3D-SIM, for comparison with the Zeiss LSM 800/Airyscan confocal image stacks and to confirm that we were not missing foci. WT cells imaged with 3D-SIM had a mean 400.6 ± 14.74 EdU foci (n = 10) while those imaged with the confocal microscope had a mean of 412.1 ± 9.38 EdU foci (n = 35). Hence, no significant difference was observed between both imaging systems (unpaired t-test; p = 0.55) (Figure 4).

**Figure 4.**
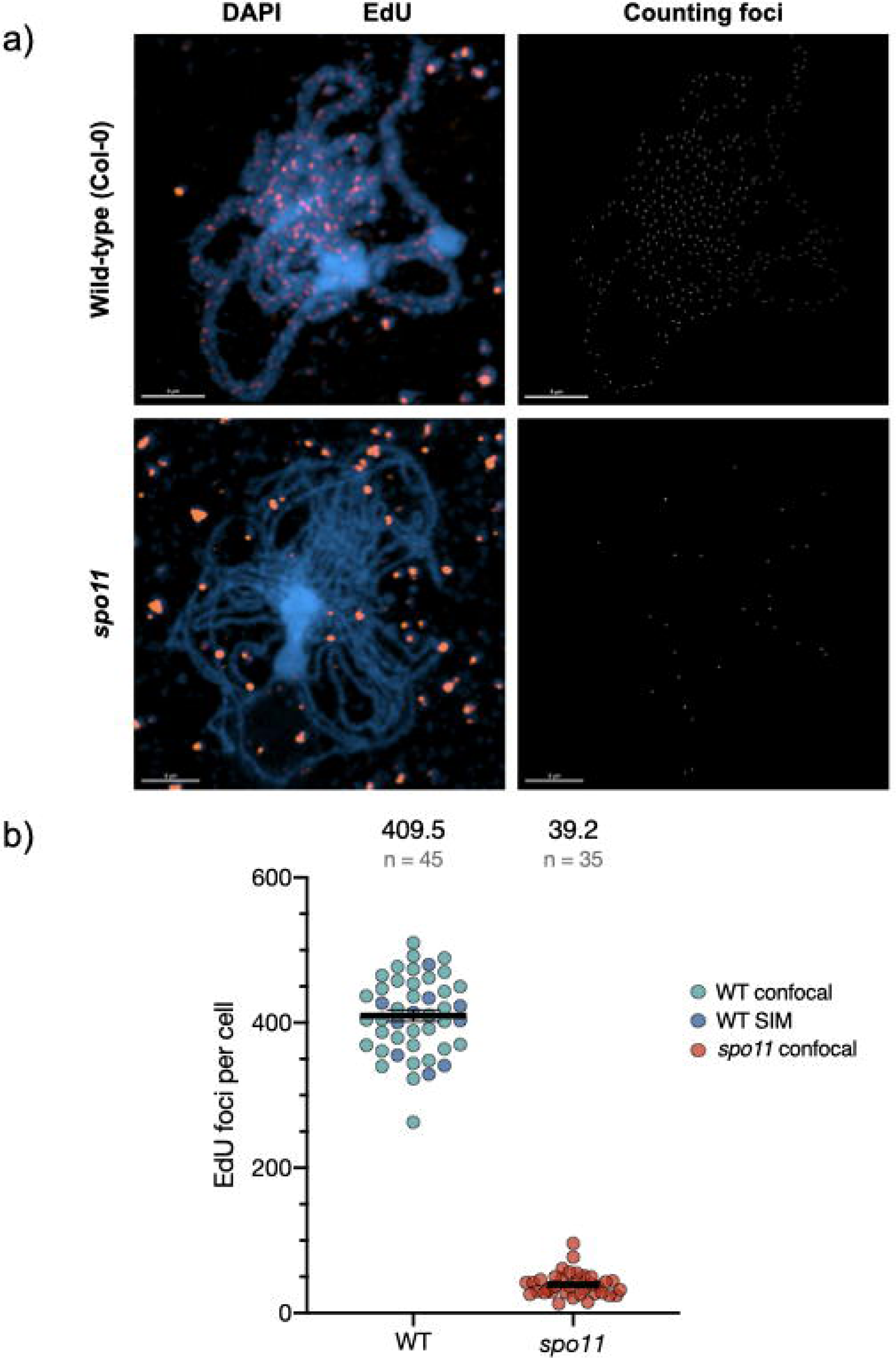
Concordance of numbers of SPO11-dependent, meiotic DNA synthesis foci and estimated DSB numbers. a) Representative images of wild-type pachytene and *spo11-1* pachytene-like meiotic stages. The merged DAPI (blue) plus EdU (orange) images are accompanied by the computer-generated images of the scored EdU foci (white, right panels). EdU foci colocalising with chromosome fibres were retained for counting and mean intensity used to exclude the much-brighter organellar foci (see text). The Images taken with the confocal microscope with Airyscan module. 3µm scale bars are shown at the bottom left of each image. b) Numbers of EdU foci on the chromosomes quantified with Imaris software show clearly the SPO11-dependence of these DNA synthesis tracts and that their numbers correspond to expectations from DSB counting in PMC. The mean number of foci/nucleus and number of nuclei counted are shown above the graph.

In *spo11-1* plants, 39.2 ± 2.83 EdU foci per cell (mean ± SEM; n = 35 cells; Figures 3, 4) were detected in zygotene/pachytene nuclei. Thus, some EdU foci were detected in *spo11-1* meioses, but in significantly lower numbers and, in general, considerably smaller and weaker than the foci in the WT. A similar signature could be observed in *spo11-1* cells at subsequent meiotic stages (Figure 3, diakinesis to Telophase II), with some EdU foci detected but in much lower numbers. This ∼10-fold reduction in mean numbers of foci with respect to the WT is highly significant (unpaired t-test; p < 0.0001) and confirms the SPO11-dependence of the great majority of the observed pachytene EdU foci. That the expected S-phase EdU labelling was observed in early prophase I *spo11-1* PMC nuclei (Figure 3, early PI, mid PI, second row) and organellar EdU labelling was comparable in *spo11-1* and WT meioses (Figures 2, 3), confirms that the EdU labelling pulse worked as expected in both WT and *spo11-1* plants.

In interpreting these numbers it is important to note that although Arabidopsis *spo11* meiosis does complete and produce (mostly inviable) pollen, chromosome pairing and synapsis are very severely reduced in *spo11* meiosis and the zygotene/pachytene stages are only identifiable by their characteristic chromosome condensation and their temporal positioning [37]. Thus, *spo11-1* PMC at leptotene can be differentiated from the zygotene-like state by the condensation of the chromosomes (thickness of the fibres) and the clustering of DAPI-dense regions. Also, *spo11-1* zygotene/pachytene-like cells that do not show S-phase EdU labelling are in a more advanced state that those that do show it, as they would have completed S-phase at the beginning of the pulse (Figure 3a, mid PI, third row vs second row). Identification of later meiotic stages in *spo11-1* meioses presents no difficulties (Figure 3).

### Prophase I DNA synthesis patterns mirror meiotic recombination features

Meiotic recombination models predict different patterns of DNA synthesis tracts depending on how recombination intermediates are formed and resolved. Non-crossover events generated by either synthesis-dependent strand annealing (SDSA) or dHJ dissolution recombination, both expected to produce DNA synthesis tracts in only one of the two implicated chromatids and thus in only one of the homologous chromosomes in interhomologue recombination. On the other hand, dHJ resolution resulting in either crossover or non-crossover products would generate DNA synthesis tracts in both participating chromatids - and both chromosomes in interhomologue events.

The expected patterns of resulting EdU foci are shown schematically in Figure 5a and examples of segments of the synaptonemal complex (SC) from SIM pachytene micrographs are shown in Figure 5b. The paired, homologous chromosomes are clearly distinguishable in these images, permitting a first analysis of different patterns of EdU foci. No obvious bias is visible in the disposition of Individual EdU foci in these images. Foci are seen at both the internal (facing the homologous chromosome) and external faces of the paired chromosomes in the SC fibre, as well as in the middle of the SC (Figure 5).

**Figure 5.**
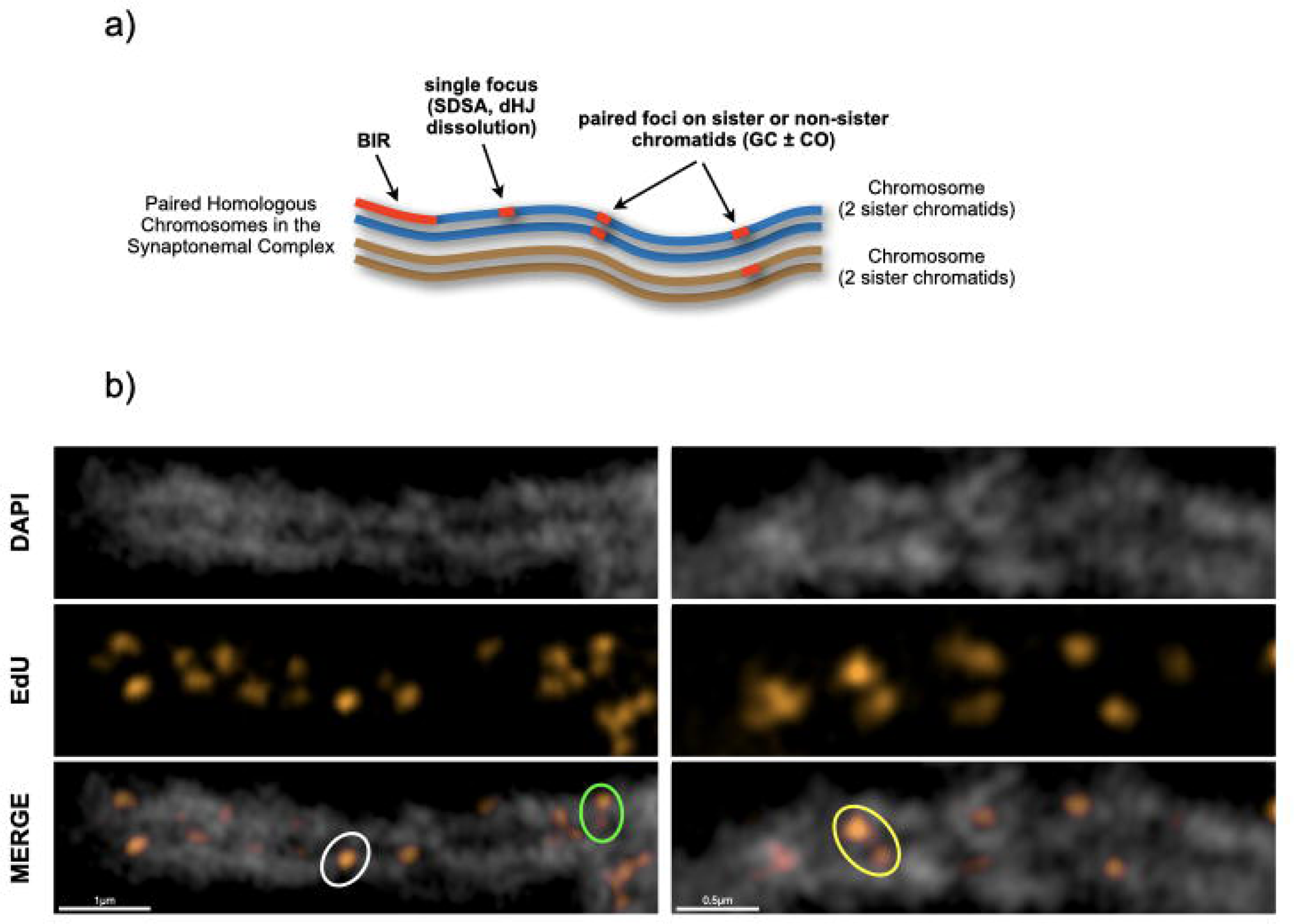
Ultrastructure of SPO11-dependent, meiotic DNA synthesis foci on the synaptonemal complex (SC) during pachytene. a) Expected configurations of DNA synthesis tracts resulting from different mechanisms of homologous recombinational repair of meiotic DSB: Synthesis-Dependent Strand Annealing (SDSA), double-Holliday Junction (dHJ), Gene Conversion ± Crossover (GC±CO) and Break-Induced Replication (BIR). b) Examples of segments of pachytene SC. Isolated individual foci (one SC lateral axis, white circle) and pairs of foci on one homologue (one SC lateral axis, green circle) or the two homologues (both SC lateral axes, yellow circle) are highlighted. The images represent a single slice of 3D-SIM image stacks. 0.5µm scale bars are shown at the bottom left of each image.

Differences can however be seen when EdU foci are analysed with respect to adjacent foci, with three clear configurations of pairs of foci, examples of which are circled on the images shown in Figure 5b:

Class (1) isolated individual foci located on one of the two homologues (one SC lateral axis; Fig. 5b, white circle).
Class (2) “pairs” of foci on one of the two homologues (one SC lateral axis; Fig. 5b, green circle).
Class (3) “pairs” of foci on the two homologues (both SC lateral axes; Fig. 5b, yellow circle).

Further examples of these patterns can be found in the pachytene images of Figure S2.

Finally, we note the absence of the long labelling tracts expected from BIR recombination, implying that such events are either rare or “short”, or both, in Arabidopsis meiosis.

### Prophase I DNA synthesis colocalize with class I crossovers

As the SPO11-dependent, meiotic prophase I DNA synthesis foci result from recombinational repair of meiotic DSB, genetic crossovers should co-localize with a subset of these EdU foci. Class I CO are detectable cytologically in Arabidopsis by the immunolocalization of proteins such as MLH1.

Combining labelling of EdU tracts with immunolocalization of MLH1 (Figure 6) gave 7.86 ± 0.24 (mean ± SEM; n = 29) MLH1 foci per WT pachytene, as expected [38]. 95.6% (n=228) of all MLH1 foci colocalized with an EdU focus and the remaining 4.4% were adjacent to one. Although this conclusion must be tempered due to the high number of EdU foci in these images, Arabidopsis Class I CO are thus physically associated with the prophase I DNA synthesis tracts at this level of resolution.

**Figure 6.**
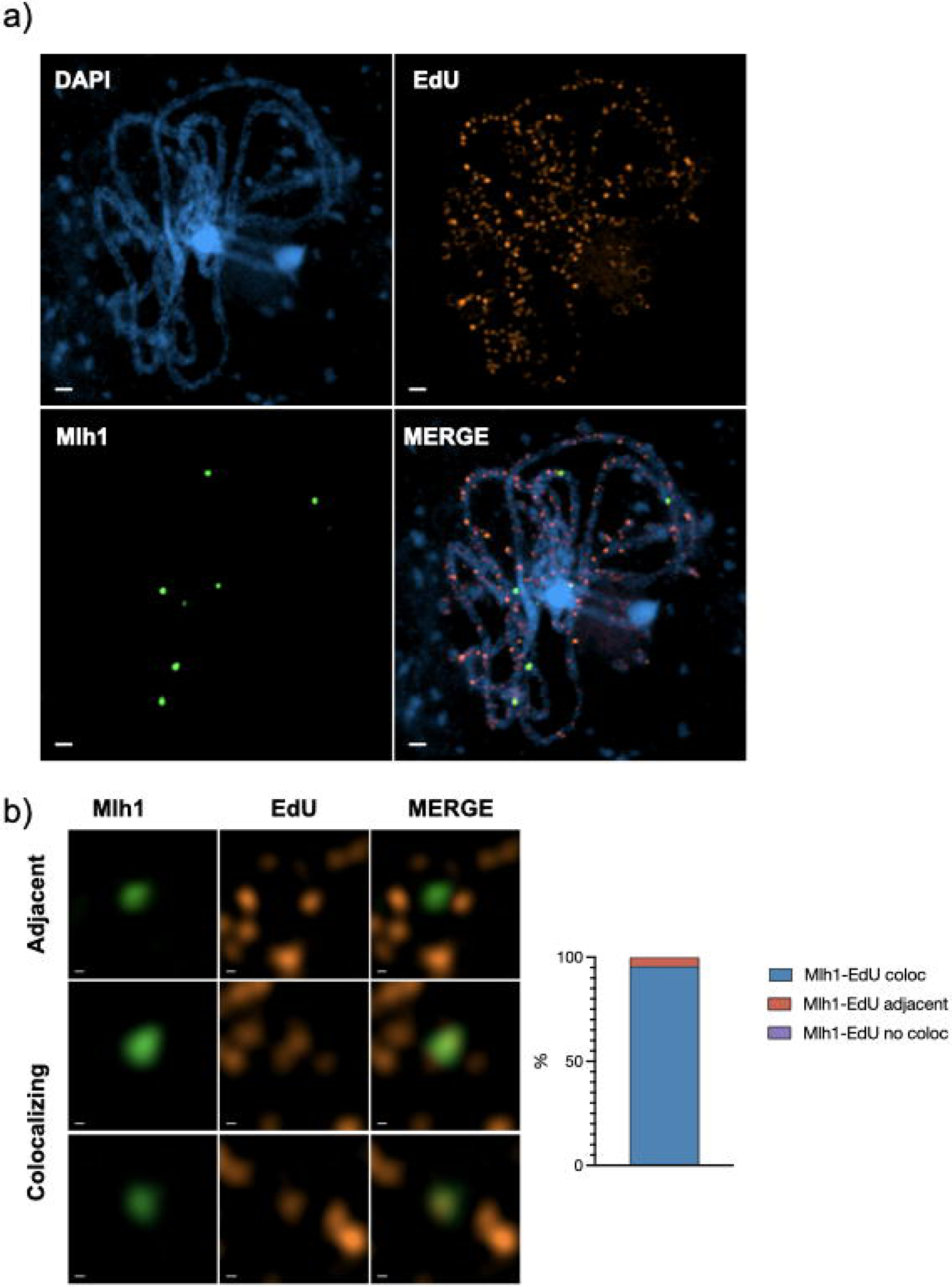
Class I crossovers (CO) colocalize with prophase I DNA synthesis tracts. a) Wild-type pachytene cell showing DAPI staining (blue), prophase I DNA synthesis tracts (EdU, orange) and Class 1 CO (MLH1, green). **b)** Examples of colocalising and adjacent MLH1 and EdU foci. The quantification (right) shows that 95.6% of MLH1 foci co-localize with EdU tracts, and the remainder are adjacent to one. a) is a confocal image and b) a single slice of a 3D-SIM image stack. Scale bars are 1µm in (a) and 0.1µm in (b).

## Discussion

### Identification of meiotic DNA repair-associated DNA synthesis tracts

We present here the labelling and characterisation of SPO11-dependent, meiotic prophase I DNA synthesis tracts through incorporation of the thymidine analogue EdU in pollen mother cells of Arabidopsis plants. By labelling DNA synthesis associated with the repair of meiotic DSB, this approach holds great promise for the future understanding of recombination patterns and mechanisms, notably for studies of NCO and sister-chromatid exchange (SCE) recombination. For practical reasons, molecular and genetic studies of meiotic recombination have essentially been focussed on events implicating the transfer of DNA sequence polymorphisms between the two recombining DNA molecules. The necessary presence of such polymorphism has, for instance, complicated significantly the analysis of the roles of meiotic SCE [39–46] and while EdU labelling has recently been applied to analysis of meiotic sister-chromatid CO in *C. elegans* (Almanzar *et al*, 2021; Toraason *et al*, 2021; Billmyre & Hughes, 2021), this approach does not appear adapted to the study of NCO.

Detection of NCO events in Arabidopsis has been elusive. Mapping NCO genome-wide by sequencing Arabidopsis tetrads has estimated NCO numbers to be of the same order as CO, ∼10 events per meiosis, with gene conversion tracts ranging between a few tens to a few hundreds of bp [19,20,50,51]. Notwithstanding this (surprisingly) low number of NCO, numbers of CO are in agreement with chiasmata counting and immunolocalisation of CO markers. Thus, given the 200-300 DSB per meiosis in the these cells, the repair of at least 90% (and very possibly >98% [52] of meiotic DSB in Arabidopsis is not accessible to analysis with current methods. In this work, we present the development of meiotic prophase DNA synthesis labelling as a means to fully characterise meiotic DSB-repair in Arabidopsis meiocytes.

Initial tests showed that incubating inflorescence stems for 24 hours in EdU solution resulted in two clear sub-populations of meiocytes, distinguishable by their meiotic stage and pattern of EdU labelling. The first consists of late G2 + early prophase I cells (leptotene, zygotene) whose chromosomes show very strong EdU signals, either genome-wide or in large patches. These patterns are consistent with those expected from pre-meiotic S-phase replicative EdU incorporation:

I) This extensive EdU labelling is observed in meioses of both wild-type and *spo11* mutant plants.
II) The extensive and intense fluorescent signals are similar to those observed in previous reports of EdU/BrdU incorporation during pre-meiotic replication [11–13,29].
III) Extrapolating from published meiotic timelines and our own studies, PMC in pre-meiotic S-phase at the beginning of the pulse are expected to advance at most to leptotene-zygotene during the 24 hour EdU pulse (see Figure 1).
IV) The signal in partially labelled early prophase I cells corresponds principally to late-replicating, DAPI-dense, heterochromatic regions, in accordance to expectations for cells in late S-phase at the beginning of the EdU pulse.

The second population of EdU-labelled meiocytes includes cells at meiotic stages from pachytene to the end of meiosis (tetrads), whose chromosomes show much fainter, discrete EdU labelling. Clearly resolving into individual foci at the stages in which the chromosomes are less condensed, these foci group into “patches” as chromosome condensation proceeds post-prophase I. These foci have a number of characteristics which permit their identification with confidence as meiotic DSB repair-associated DNA synthesis tracts:

I) These discrete EdU-label foci are SPO11-dependent, analogously to earlier observations in yeast [9], confirming with high confidence that they result from EdU incorporation at DNA synthesis tracts associated to meiotic DSB repair. Some EdU foci are observed in *spo11-1* meioses, although at considerably lower numbers than in the wild type (39 versus 410 per meiosis respectively). While some of these may come from mistaken assignation of organellar DNA foci, we note that some crossing-over is detected in Arabidopsis *spo11* plants, RAD51 foci are detected in Arabidopsis *spo11* meiosis and DMC1 foci in rice meioses [37,53–55]. Furthermore, given that both meiotic Arabidopsis SPO11 proteins are needed for normal meiotic DSB induction, some leakage may occur in plants lacking one or the two SPO11 proteins [54,56–59]. Alternatively, they may correspond to SPO11-independent breakage of mechanical, chemical or enzymatic origin. If so, that they do produce DNA neo-synthesis tracts argues for their repair by HR rather than End Joining.
II) The SPO11-dependent foci are seen at later meiotic stages than those with replicative labelling and these nuclei had thus completed S-phase prior to the beginning of the pulse. Given the known timing of meiosis in these cells, meiotic DSB repair DNA synthesis will have occurred during the EdU pulse in these cells [9,13,14,38,60]. We note that while present in considerably lower numbers than in the wild type (39 versus 410 per meiosis, respectively), there are some EdU foci in *spo11-1* meioses.
III) The discrete EdU foci observed at pachytene and later stages are both much less intense and smaller in size than the generalised labelling in leptotene-zygotene nuclei. This is expected given the predicted size of DNA synthesis tracts associated to recombination events (few bases to some kilobases of DNA), with respect to replicative labelling (megabases to the full genome).
IV) The numbers of EdU foci measured at pachytene concord with published estimations of numbers of meiotic DSB and HR intermediates in Arabidopsis. A mean of 409.5 ± 7.96 (mean ± SEM; n = 45 meioses) DNA repair-associated DNA synthesis foci were observed in WT pachytenes. This number is a little higher than estimations of meiotic DSB numbers through immunolocalisation of early recombination proteins in Arabidopsis (RAD51 in most cases). Reported numbers of RAD51 foci per Arabidopsis meiosis range from 90 to 250 per meiosis, and compare well with a recent report of 239 ± 30 (mean ± SD) prophase I SPO11-1 foci [56]. Given the limitations of immunodetection and that the recruitment/release of these proteins at DSB/recombination sites is dynamic, these numbers presumably under-estimate the true values. Notably, a recent report using a STED super-resolution microscope found more than 1000 RAD51 foci per PMC meiosis [52]. Thus, the mean of 409.5 DNA repair-associated synthesis tracts per meiosis reported here is probably closer to the true value than that of protein-immunodetection approaches. It is of course necessary to take into account the fact that resolution of dHJ recombination intermediates is expected to result in patches of neo-synthesized DNA on both implicated chromatids, resulting in two foci from a single recombination event. Such events are however estimated to represent ∼5% or less of meiotic DSB repair events in Arabidopsis and so are not expected to impact significantly the estimations of total events.
V) DAPI-dense heterochromatic, centromeric regions have a visibly lower density of SPO11-dependent EdU foci in pachytene/diplotene and beyond (particularly noticeable at diakinesis/MI). This concords with SPO11-oligo mapping of meiotic DSB patterns in Arabidopsis PMC, which shows that centromeric regions have a much lower density of DSB than the rest of the chromosome [34,36].
VI) Combining labelling of EdU tracts with immunolocalization of MLH1 confirms that, as expected, Class I CO are physically associated with the prophase I DSB repair DNA synthesis tracts (Figure 6).

### Meiotic DSB repair-associated DNA synthesis tracts mirror meiotic recombination features

As presented in Figure 7, different mechanisms of recombinational repair of meiotic DSB will result in different patterns of DNA synthesis tracts associated to an HR event. The most obvious difference is that DNA synthesis is expected to be found on only the recipient chromatid, which carried the initiating DSB (Figure 7f), or on both the recipient and donor chromatids (Figure 7g, h). SDSA and dHJ dissolution mechanisms would lea d to the former and dHJ resolution to the latter. A special case is the BIR mechanism, which is expected to result in a long DNA synthesis tract, potentially extending all the way to the end of the recipient chromatid (Figure 7c).

**Figure 7.**
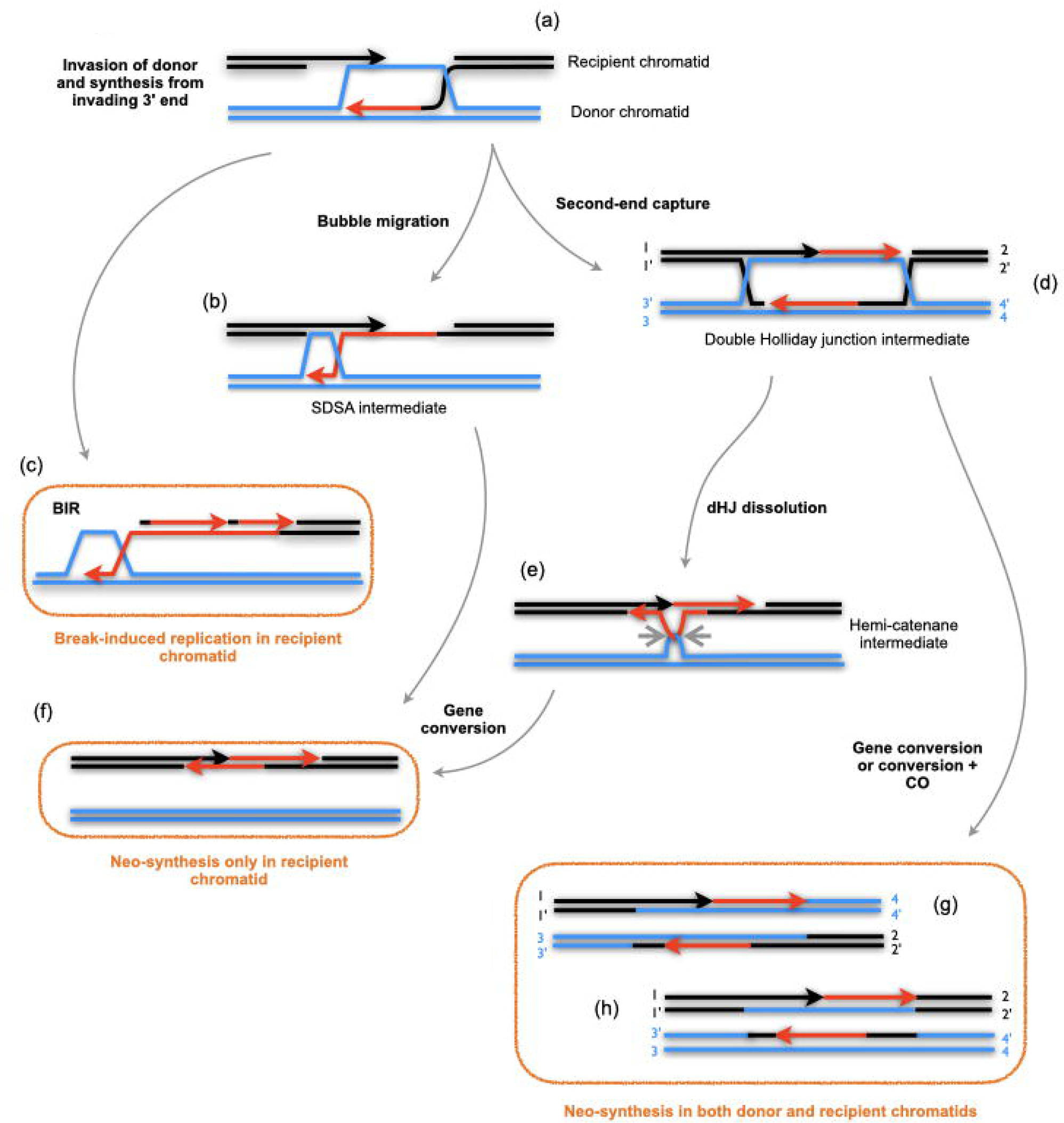
Recombination mechanisms and DNA synthesis tracts. Following resection and invasion of the donor chromatid (a), DNA neo-synthesis tracts resulting from repair of DSB are expected to be found only on the recipient chromatid in gene conversions (f) resulting from SDSA (b) or dHJ dissolution (e) pathways. A special case of this is the BIR pathway (c), which will result in a long synthesis tract, potentially extending all the way to the end of the recipient chromatid. DNA synthesis tracts are expected to be found on both the recipient and donor chromatids in gene conversions, associated (g) or not (h) to a CO, resulting from resolution of the dHJ intermediate (d).

Visual inspection of the pachytene images clearly shows different configurations of the EdU foci, which we have arbitrarily divided into 3 classes suggestive of the DNA synthesis patterns expected at sites of recombination:

Class (1) isolated individual foci located on one of the two homologues (one lateral axis) (Fig. 5b, Fig. S2). Only the initially broken chromatid (recipient) will bear (EdU-substituted) neosynthesized DNA tracts in repair via SDSA (Fig. 7a, b, f) and dHJ dissolution (Fig. 7a, d, e, f) recombination.

Resolution of double Holiday junction (dHJ) intermediates (Fig7d) can give rise to gene conversions, accompanied (Fig. 7g) or not (Fig. 7h) by a CO. In both cases, this can lead to the presence of neo-synthesized DNA on both the donor and recipient chromatids. These events would thus be expected to result in paired DNA synthesis tracts on the two participating chromatids: either on one chromosome (Class (2) - one lateral axis) for sister chromatid events, or on both homologues (Class (3) - both lateral axes).

Class (2) are thus “pairs” of foci on one of the two homologues (one SC lateral axis), which could result from recombination between sister chromatids through the dHJ resolution pathway.

Class (3) are “pairs” of foci on the two homologues (both lateral axes), which could result from recombination between homologues through the dHJ resolution pathway.

We find no evidence for long DNA synthesis tracts indicative of BIR recombination in our images. Thus, either BIR replication tracts are not distinguishably longer than other DSB repair-DNA synthesis, or the BIR mechanism is a minor (or non-) contributor to DSB repair in Arabidopsis meiosis. In this context, we note that an earlier effort to detect mitotic BIR repair in bleomycin-treated *Vicia faba* root tips described asymetric interstitial G2-phase EdU labelling of interstitial tandem-repeat regions, although no extended labelling tracts out to the end of the chromatid were observed [61].

Inference of the recombination pathways underlying the 3 classes of foci of course remains speculative in the absence of molecular analyses of the EdU tracts themselves. FISH on extended DNA fibres was used in an earlier study in yeast to determine the lengths and positioning of BrdU-substituted DNA synthesis tracts with respect to DSB hotspots [9]. They observed two main classes of tracts: tracts that extended from the DSB towards only one side of the DSB and tracts that extended on both sides. These two classes correspond to the tracts expected in single DNA molecules that took part in CO events in the first case and NCO events via SDSA in the second, which were further validated studying their dynamics in recombination mutants. We have previously carried out DNA combing plus fibre FISH in Arabidopsis [62], but the lack of strong meiotic DSB hotspots with a precisely localised DSB site complicates the possible use of this approach in Arabidopsis meiosis. Rather, we are actively developing an approach based on the direct detection of EdU-substituted DNA tracts by Nanopore sequencing. It is hoped that this approach will permit unambiguous characterisation of the principal meiotic recombination products and underlying mechanisms in Arabidopsis, as well as of more complex events.

## Materials and Methods

### Plant material and growing conditions

Standard conditions were used for growth of wild-type *A. thaliana Col-0* and *spo11*-1-2 [37] plants. Seeds were sown on soil substrate (Klasmann-Deilmann GmbH TS3 FIN 416GF, Geeste, Germany), stratified for 2-4 days at 4°C and grown in climate chambers at 23°C and 60% relative humidity, under a daily cycle of 16 hours of light (110-140 μmol m^−2^ s^−1^) and 8 hours of darkness.

### Meiotic chromosome preparation by spreading

Meiosis was analysed following the chromosome spreading method of Ross et al. [63], with minor modifications. Inflorescences were collected in 3:1 ethanol:glacial acetic acid fixative solution and left overnight at room temperature. The fixative was then substituted with fresh fixative solution and the tubes were transferred to 4°C. This step was repeated the following 2-3 days. At this point, fixed inflorescences can be stored at 4°C for further use or prepared for microscopy.

A number of fixed inflorescences were selected, deposited in a glass dissection well with 3:1 fixative solution and dissected under a stereomicroscope into individual flower buds, discarding open flowers and buds with pollen (yellow anthers). The flower buds were then washed 3 x 2 minutes in citrate buffer and the buffer replaced with 500 μl of digestion enzyme mix (0.3% w/v cellulase from *Trichoderma* sp. (Sigma-Aldrich), 0.3% w/v pectolyase from *Aspergillus japonicus* (Sigma) and 0.3% w/v cytohelicase from *Helix pomatia* (Duchefa Biochemie) in citrate buffer). The flower buds were incubated in the digestion mix for 2 hours at 37°C and the digestion stopped by adding a greater volume of ice-cold citrate buffer. The well was placed under the stereomicroscope and a single flower bud with a small volume of liquid was transferred to a slide with a Pasteur pipette. The flower bud was macerated with a needle, a 10μl drop of ice-cold 60% glacial acetic acid was added and the slide incubated for one minute on a hot plate at 45°C while gently moving the drop with a flat needle. After the incubation, an extra 10 μl of ice-cold 60% glacial acetic acid were added, followed by 100 μl of 3:1 fixative - first circling and flooding the drop and then mixing with the drop by vigorously pipetting the last microliters. The liquid was drained by tilting the slide, which was then further washed with another 100 μl of 3:1 fixative, drained and left to air dry at room temperature.

To visualize the chromosomes, the slides were stained in Vectashield antifade mounting medium with 1.5 μg/ml DAPI (# H-1200, Vector Laboratories, Burlingame, CA, USA), by putting a drop on a coverslip which was then placed on a slide and gently squashed by thumb pressure. Spreads were analysed with a Zeiss AxioImager Z1 microscope with a 100×/1.40 Oil Plan-Apochromat objective and the Zeiss ZEN2Blue software (Carl Zeiss GmbH)in the channel for blue fluorescence (Zeiss #49 HE filter).

### EdU labelling and cytological detection

Among the different thymidine analogues commonly used to label *in vivo* DNA synthesis, 5-Ethynyl-2’-deoxyuridine (EdU) has a number of advantages and has been preferred by the Arabidopsis community in recent studies labelling DNA synthesis [12,29,30,64–67]. EdU can be detected in intact double-strand DNA and this antibody-free technique simplifies the protocol and offers better reproducibility.

EdU (#Ab146186, Abcam, UK) working solutions were prepared at 10mM in 1×PBS buffer. Arabidopsis stems with apical inflorescences were submerged in water to avoid the introduction of air bubbles (submerging only the stem segment to be cut, without wetting the inflorescences) and cut obliquely 4-5cm below the inflorescence. Care was taken to make the cleanest cut possible in order to avoid crushing vascular tissues. Open flowers and big leaves (if present) were removed to avoid crowding the tube and minimize the risk of exhausting the solution due to excessive transpiration. The inflorescences were incubated in 800μl of 10mM EdU in a 2ml eppendorf tube for 24 hours, under the same growing conditions as the plants. To avoid tight packing, a maximum of four or five inflorescence stems were incubated in one tube. After 24 hours the inflorescences were removed from the EdU solution and fixed in 3:1 ethanol:glacial acetic acid solution.

Chromosome spreads were prepared as previously described and visualised under the microscope to identify those slides with meiocytes at the desired meiotic stages. The slides were first incubated in 4T (4×SSC, 0.5% v/v Tween 20) in a Coplin jar until the coverslips detached, followed by 30’ without coverslips in fresh 4T. The Click-iT reaction mix (Invitrogen Click-iT plus Alexa-fluor 647 kit, Thermo-Fisher Scientific) to label EdU with the fluorochrome was prepared following the manufacturer’s instructions. Before labelling, the slides were washed for 5 minutes with 1×PBS. A 30μl drop of Click-iT reagent was placed on a 32×24mm piece of parafilm and the inverted slides were placed on top of the mix on the parafilm. The slides were incubated with the mix for 30 minutes at 37°C in a dark moist chamber, washed for 5 minutes with 1×PBS in a Coplin jar in the dark and mounted in Vectashield + DAPI for microscopy.

Pictures were taken using a Zeiss LSM800 confocal microscope applying the Airyscan module and selecting the presets for each specific dye. EdU foci were quantified semi-automatically using the IMARIS software *Spots* tool (Bitplane AG, Oxford Instruments, UK), adding intensity filters for both channels to specifically selected visually detectable EdU foci that co-localised with the DAPI fluorescence of the chromosomes. To analyze the chromatin and EdU signal ultrastructure, we applied super-resolution spatial structured illumination microscopy (3D-SIM) using a 63×/1.40 Oil Plan-Apochromat objective of an Elyra PS.1 microscope system and the ZENBlack software (Carl Zeiss GmbH) [68]. Maximum intensity projections were calculated from 3D-SIM image stacks. Zoom-in sections were presented as single slices to indicate the subnuclear chromatin structures at the super-resolution level.

### Detection of MLH1 and EdU

For the detection of MLH1 and neo-synthesized DNA tracts with incorporated EdU, the protocol for the immunolocalization of MLH1 [69] was adapted to include the Click-iT EdU labelling. Chromosome spreads were prepared as previously described and visualized under the microscope to identify those slides with meiocytes at the desired stages. The slides were incubated in a Coplin jar with 100% ethanol to detach the coverslip and washed with 1×PBST (0.1% Triton X-100 in 1×PBS. Another glass staining jar was filled with tri-sodium citrate buffer pH7 and heated to boiling point in a microwave. The slides were placed in the heated tri-sodium citrate solution for 45 seconds, after which they were returned to room temperature 1×PBST.

The rabbit anti-MLH1 primary antibodies (kindly provided by Mathilde Grelon, INRAe Versailles [38]) was prepared at 1/200 in 1×PBST + 1% BSA. A 45μl drop was placed on a 32×24mm piece of parafilm and the inverted slide place on this. The slides were incubated for 2 days at 4°C in a dark moist chamber, after which they were washed 3 x 5 minutes with 1×PBST. The secondary antibodies (goat anti-rabbit Alexa fluor 488) was prepared at 1/100 1×in PBST + 1% BSA, and 45μl were used per slide. Slides were incubated 30 minutes at 37°C in a dark moist chamber, after which they were washed 2 x 5 minutes in 1×PBST, followed by one wash with 1×PBS. The Click-iT reaction was then performed as described above and the slide was mounted in Vectashield + DAPI mounting medium for microscopy. Counting of MLH1 foci and co-localization with EdU foci was carried out by visual inspection of images taken using the Zeiss LSM800 confocal microscope plus the Airyscan module.

## Acknowledgements

We thank the members of the recombination group and especially Olivier Da Ines, Jérémy Verbeke and Floriane Chéron for their help and suggestions. Mathilde Grelon kindly sent us the anti-MLH1 antiserum.

This work was supported by a European Marie Sklodowska Curie Actions Innovative Training Network grant (EU H2020-MSCA-ITN-2017-765212) to CIW, the Centre National de Recherche Scientifique (CNRS), the Institut National de la Santé et de la Recherche Médicale (Inserm) and the Université Clermont Auvergne. The funders had no role in study design, data collection and analysis, decision to publish, or preparation of the manuscript.

## Supplemental Figure Legends

**Figure S1.**
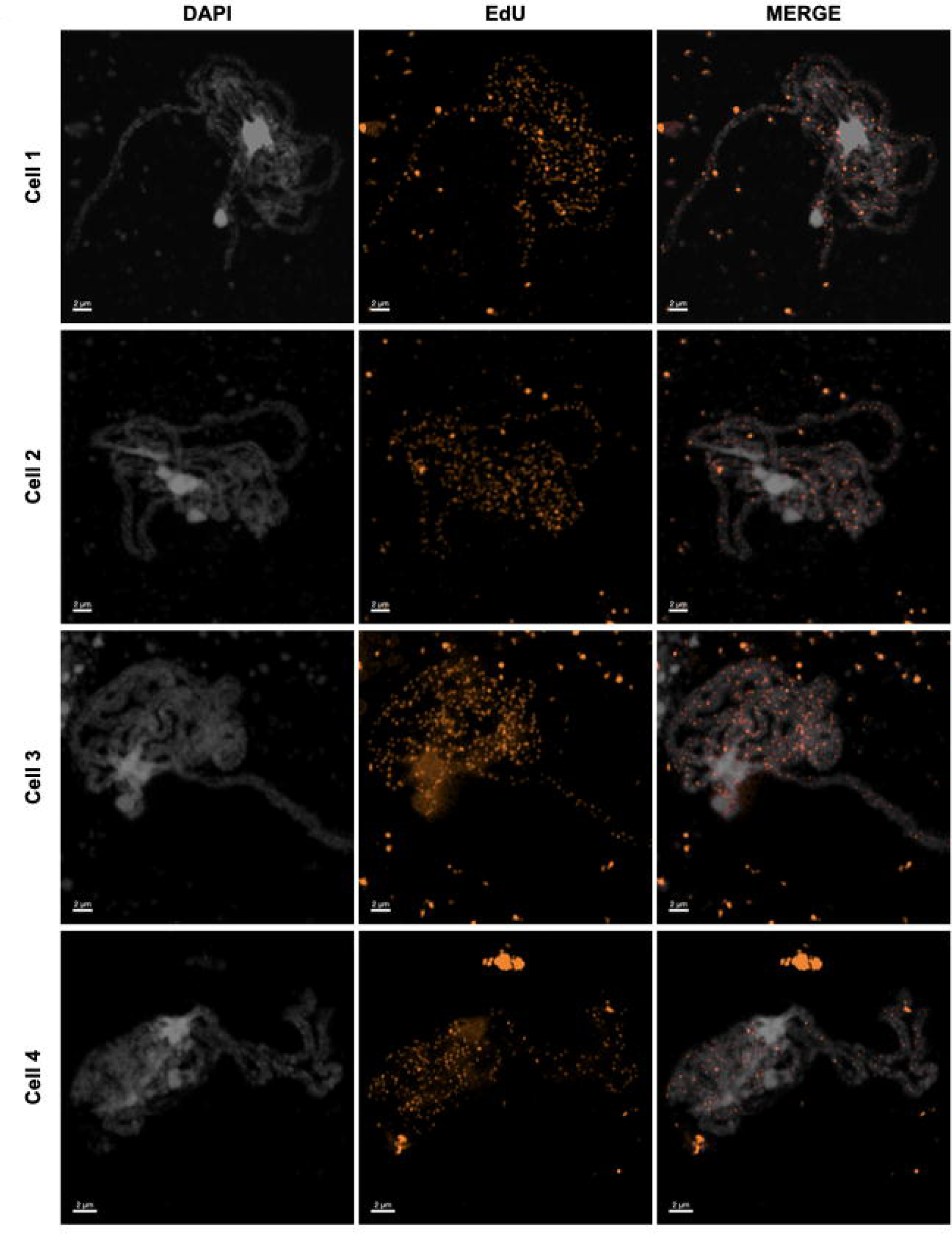
Discrete foci of EdU substituted DNA are visible in pachytene chromosomes of Arabidopsis PMC. Examples of prophase I DNA labelling in pachtene nuclei of wildtype PMC. Each row is the same nucleus, imaged with DAPI fluorescence (white, left), EdU (orange, middle) and the merged image (right). Images taken with the confocal microscope with Airyscan module. 2µm scale bars are included at the bottom left of each image.

**Figure S2.**
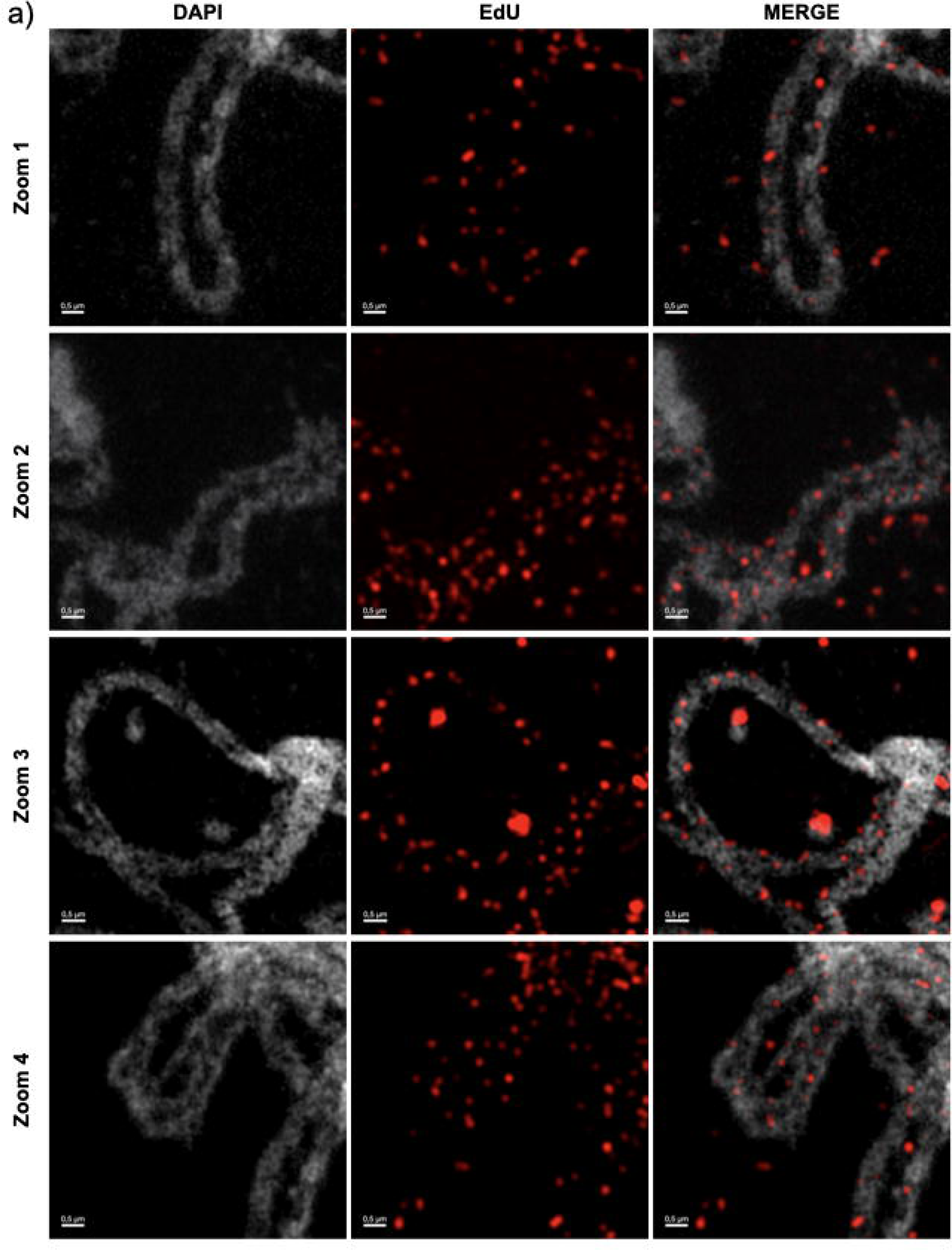
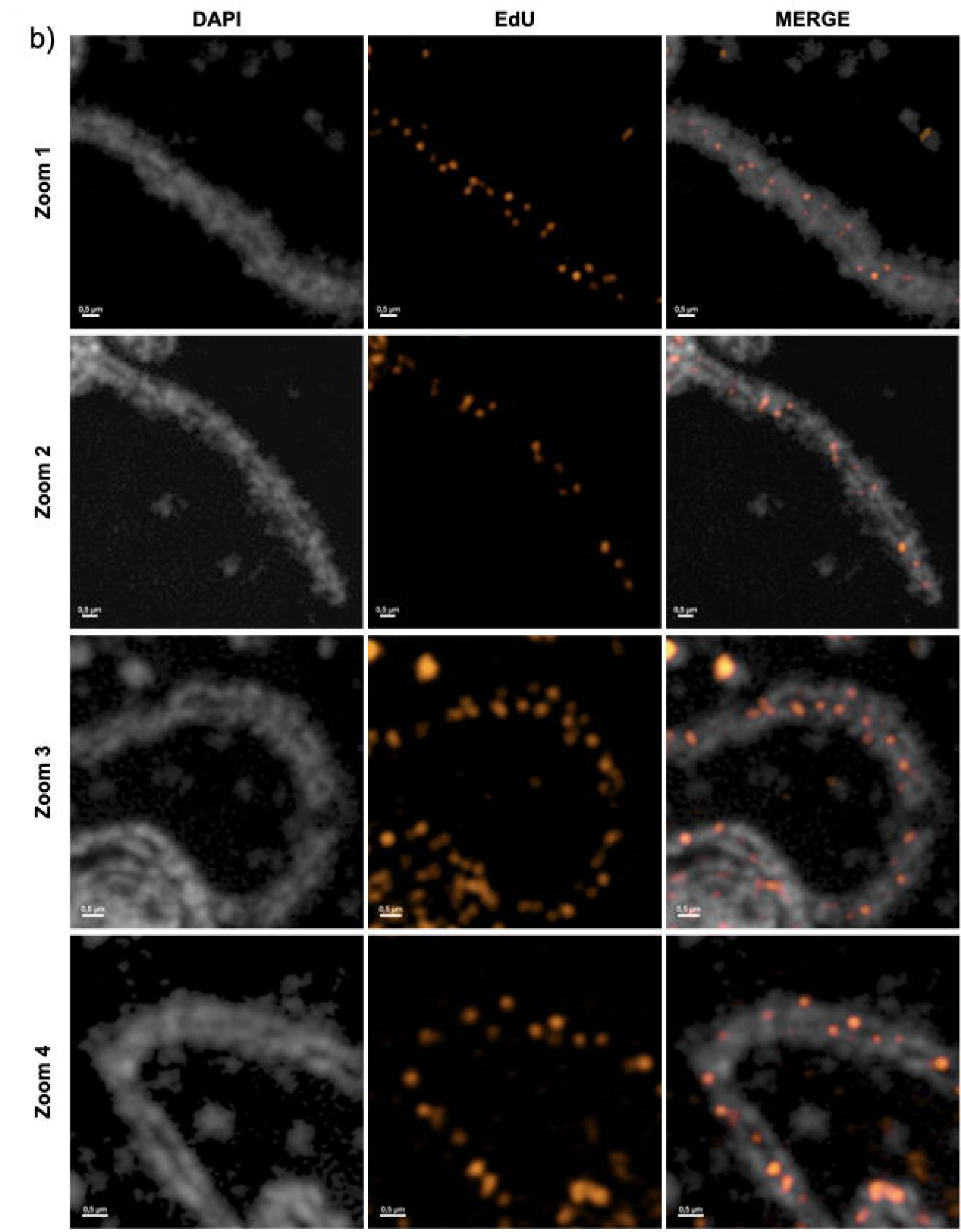
Distribution of SPO11-dependent, meiotic DNA synthesis foci on paired pachytene chromosomes from two different cells. Zoomed regions from SIM (a) and Confocal (b) images of meiotic pachytenes, showing the distribution of EdU-labelled meiotic prophase I DNA synthesis foci (a)red, b) orange) on the DAPI-stained chromosome fibres (white) of the synaptonemal complexes. 0.5µm scale bars are included at the bottom left of each image.

